# Deep learning of protein energy landscape and conformational dynamics from experimental structures in PDB

**DOI:** 10.1101/2024.06.27.600251

**Authors:** Yike Tang, Mendi Yu, Ganggang Bai, Xinjun Li, Yanyan Xu, Buyong Ma

## Abstract

Protein structure prediction has reached revolutionary levels of accuracy on single structures, implying biophysical energy function can be learned from known protein structures. However apart from single static structure, conformational distributions and dynamics often control protein biological functions. In this work, we tested a hypothesis that protein energy landscape and conformational dynamics can be learned from experimental structures in PDB and coevolution data. Towards this goal, we develop DeepConformer, a diffusion generative model for sampling protein conformation distributions from a given amino acid sequence. Despite the lack of molecular dynamics (MD) simulation data in training process, DeepConformer captured conformational flexibility and dynamics (RMSF and covariance matrix correlation) similar to MD simulation and reproduced experimentally observed conformational variations. Our study demonstrated that DeepConformer learned energy landscape can be used to efficiently explore protein conformational distribution and dynamics.

## Introduction

Accurate methods for protein structure prediction such as AlphaFold^1,2^ and later methods^3–5^ have greatly pushed forward understanding of protein function and the rational design of therapeutics^42^. However, while such methods are designed to model static experimental structures from crystallography or cryo-EM as seen in PDB, proteins adopt dynamic structural ensembles featuring conformational flexibility in solution. Understanding the dynamical properties of proteins beyond their static structure is crucial to reveal their biological function^6,7^. Therefore, the reliable sampling of structure distributions due to conformational dynamics based on protein sequence has increasing importance^8^.

The problem of predicting an experimental-level protein structure from sequence is widely considered to have been solved by AlphaFold2^2^. Since then, alternative models such as RoseTTAFold^3,5^ also reached similar accuracy. Both Alphafold2 and RoseTTAFold are based on multiple sequence alignments (MSA) input. Methods based on single sequence input and pretrained protein language model such as ESMFold^4^ have approached similar levels of performance, slightly less accurate than Alphafold2/3. All these methods are developed and trained as deterministic maps from input (sequence or MSA) to output (structure), making them suboptimal for modeling structural ensembles ^9^. MSA subsampling and clustering (i.e., varying the input) in conjunction with AlphaFold2 has recently been shown to reveal alternate conformations^10–14^, but all these studies limited these study and comparisons to existing alternative protein conformations.

Generative networks especially diffusion models ^15,16^ have been applied to biomedical problems such as protein design^17–22^, molecular docking ^23,24^, and ligand design^25,26^. Such models have displayed impressive distributional modeling. These capabilities make diffusion models compelling tools for understanding protein structural ensembles given a fixed sequence, demonstrated by recent studies of using diffusion models developed for protein structures prediction (^27,28^, EigenFold^29^, AlphaFold3^1^, PVQD ^30^).

AlphaFold3^1^ replaces the deterministic structure module in AlphaFold2 with a diffusion-based model. Like other protein structure predictions models a key limitation is that it predicts static structures as seen in the PDB, not the dynamical behaviour of biomolecular systems in solution.

DiG^31^ relies on input of MD generated conformation for training. Training generative-AI model on the across-system conformation datasets sampled from MD simulation is a natural way to let model learn protein conformational dynamics. However, the amount of such data is limited due to the computationally intensive MD simulations and MD simulation itself may have problem sample complete conformation space due to high energy barriers among different energy basins on protein energy landscape.

Liu and Bahar have shown that protein sequence evolution correlates with structural dynamics ^32^. Roney and Ovchinnikov^33^ argued that AlphaFold2 has learned biophysical energy function from known protein structures, and the learned energy function can be used to rank the quality of candidate protein structures without using any coevolution data^33^. Therefore, we hypothesized that protein energy landscape and conformational dynamics can be learned from experimental structures in PDB and coevolution data. In this work, we tested this hypothesis and developed a model named DeepConformer which only used expanded experimental protein structures, without reliance on the computationally intensive simulation data. Our study demonstrated that DeepConformer has learned protein energy landscape and can be used to efficiently explore protein conformational distribution and dynamics.

## Materials and Methods

### 1. An overview of DeepConformer

The forward and reverse process of our diffusion model are adopted from RFdiffusion^17^. Namely, we used the Cartesian coordinates of C-alpha (represented by p) and the rotation matrix of the triangle formed by [N, C-alpha and C] (represented by R) to represent the geometric position of a residue backbone.

The perturbation kernel 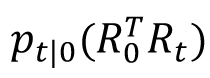 for the rotation components 𝑅_𝑡_ is considered element-wise via the isotropic Gaussian on SO(3) distribution ^17,18,34^:

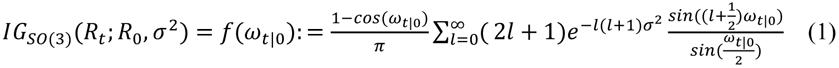

where 𝜔_𝑡|0_ = axis-angle (𝑅^𝑇^𝑅_𝑡_) is the rotation angle in radians of the rotation matrix 𝑅^𝑇^𝑅_𝑡_. On the other hand, the perturbation kernel for translation components pt is an isotropic Gaussian kernel.

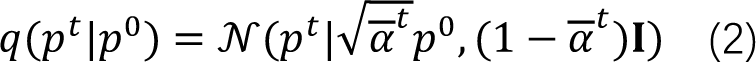

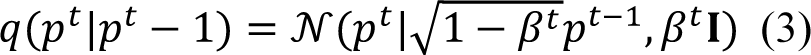

where 𝛽^𝑡^ controls the rate of diffusion and its value increases from 0 to 1 as time step goes from 0 to t, and 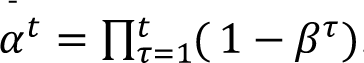. As the coordinate of an atom could be an arbitrary value, we zero-center the coordinates of C-alpha and multiply coordinates by 0.25 to make the distribution of atom coordinates roughly match the standard normal distribution.

In addition to the residue coordinates, the network is provided with node and edge features obtained by running AlphaFold on the input sequence and extracting the node and pair embeddings from the Evoformer stack.

The object of training the generative process is to recover noised backbone structure conditioned on the extracted node and pair embedding. The overall loss function is formulated to minimize the discrepancy between the ground truth structure and predicted de-noised structure:

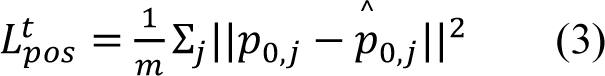

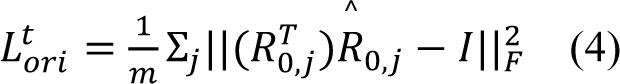

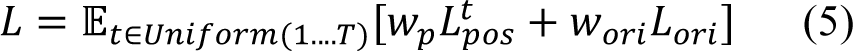

Where m is the number of residues in the protein. The training process can be described with following steps:

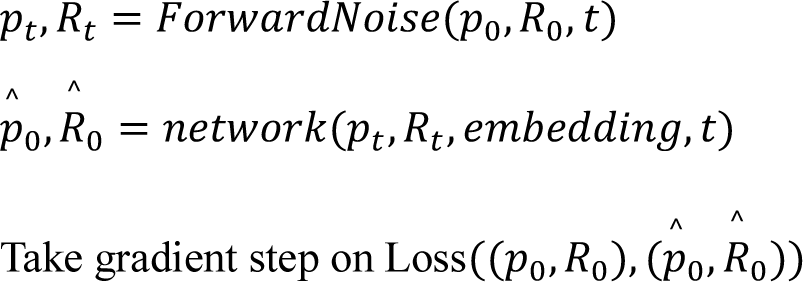

In the inference process, initial backbone structure is generated from random noise 𝑝_𝑇_ ∼ 𝒩(0, 𝐼) and 𝑅_𝑇_ ∼ 𝑈𝑛𝑖𝑓𝑜𝑟𝑚(𝑆𝑂(3)).For t=T,….,1

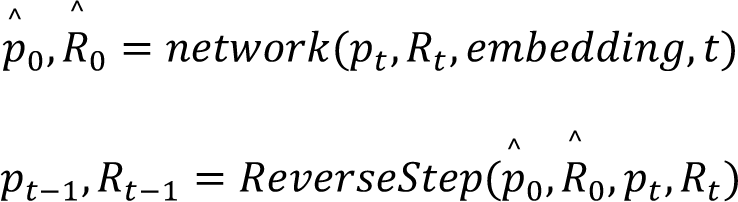

Here ReverseStep function is taken from RFdiffusion(Algorithm 2).

Our neural network is modified from the structure module of AlphaFold2 (AF2). Several previous works have adopted similar network structure for protein structure diffusion ^18,35^. The initial single embedding from AlphaFold2 is concatenate with sinusoidal encoding of the time step and then pass to a linear layer. The initial pair embedding is concatenate with sinusoidal encoding of the time step, relative residue position encoding, as well as RBF encoding of the initial pairwise distance between ||𝑝_𝑡,𝑖_ − 𝑝_𝑡,𝑗_ || and then passed to a linear layer. The composed node and edge embedding together with initial structure parameter (𝑝_𝑡_, 𝑅_𝑡_) are passed to 5 consecutive layers of neural network. Fig. 1 shows one single layer of our neural network. Spatial attention is performed with Invariant Point Attention (IPA) from AF2 which can attend to closer residues in coordinate space and capture interactions along the chain structure. The updates are SE(3)-invariant since IPA is SE(3)-invariant. We utilize fully connected graph structure where each residue attends to every other node. Updates to the node embeddings are propagated to the edges in EdgeUpdate where node embedding ℎ_𝑖_, ℎ_𝑗_ and edge embedding 𝑧_𝑖,𝑗_ are concatenated and passed to a MLP which outputs an updated edge embedding. BackboneUpdate is taken from AF2 (Algorithm 23), where a linear layer is used to predict translation and rotation updates to each frame.

**Fig 1.**
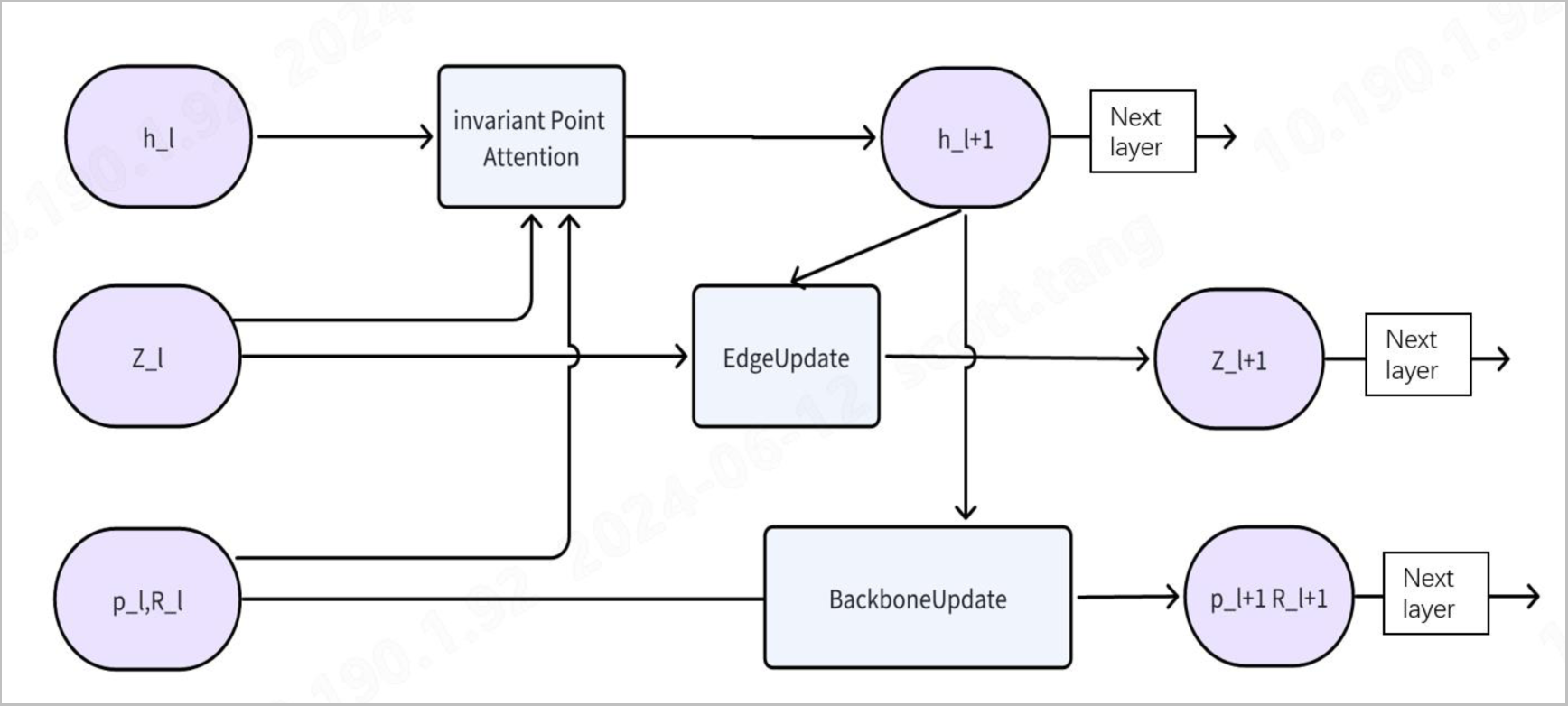
Schematic diagram of single layer neural network.

#### 1. 2. Training and Sampling Details

Three elements are introduced into Deepconformer to increase its ability to learn comprehensive energy landscape. Figure 2 illustrate the first two: (1) associate different structures for a given protein sequence. (2) Large scale masked amino acid positions (50-70%) are randomly selected to increase sequence-structure association during recovery process.

**Fig 2.**
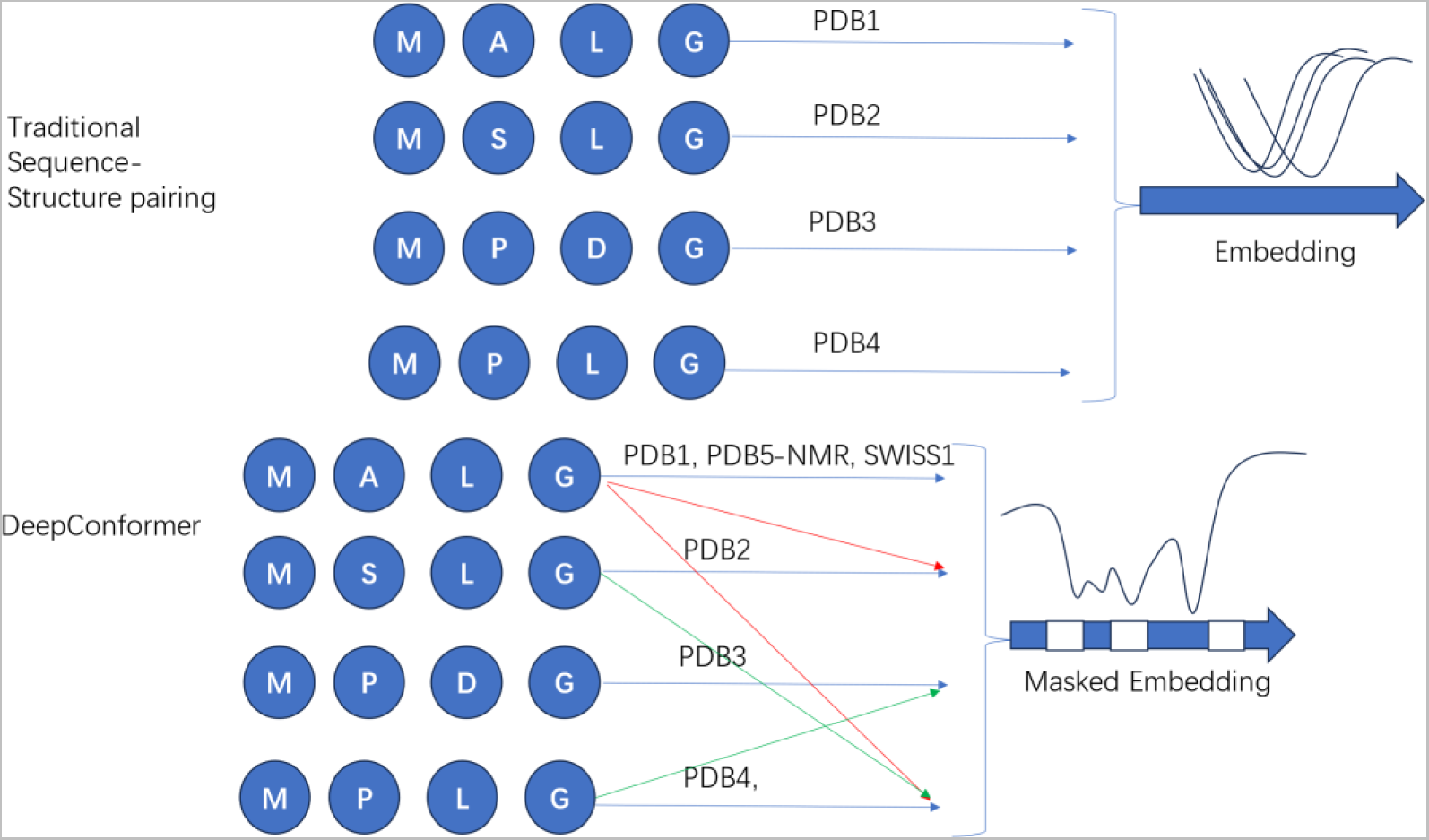
Deepconformer associates different structures on a protein energy landscape for a given protein sequence. Large scale masked amino acid positions (50-70%) are randomly selected to increase sequence-structure association. Upper: traditional pairing of protein sequence with a known protein structure. Lower panel: Deepconfomer masked energy landscape embedding.

Traditional methods trained AI models with PDB structure and corresponding amino acid sequence. Most sequences in PDB have only one or two conformations and lack conformational diversity. However, structures determined by NMR method in PDB usually have 10 to 20 conformations and therefore contain more distributional information than structures determined by X-ray or Cryo-EM. Therefore, we used all conformers in a PDB entry determined by NMR method. Secondly, we argued that two proteins with a single amino mutation share one parental energy landscape, and mutation usually shifted the energy landscape slightly. Therefore, to enhance the conformational diversity, we developed a set of techniques, we merged {sequence: structures} pairs if the difference between the two sequences is a single point mutation. A part of the sequences in PDB also have alternative conformations in Swiss-Model repository^36^ and can be added into training dataset.

The amino acid (node) mask also leads to subsequent edge mask which is associated with the masked node. Protein within the same fold class could merely share less than 30% sequence identity ^37^. Therefore, we introduced large scale amino acid positions mask (50-70%) to increase sequence-structure association. In both training and inference process we randomly selected 50% of the residues and mask their corresponding node and edge embedding (set to zero). In the inference process at each time step a different set of residues were selected.

The third element to increase the coverage of energy landscape associated with a given protein sequence is to use MSA clustering. Wayment-Steele et al. ^11^ found that by clustering MSA via input AF2 can predict distinct conformations of a single protein sequence. Inspired by their work, we developed a pipeline for proteins that have broader conformation spaces. When extract embedding from Alphafold2 we first cluster MSA via sequence identity, top 5 clusters with max sequences are selected and passed to Alphafold2 independently. Correspondingly, we obtain 5 embeddings. These embeddings are averaged, then feed to Deepconformer model. With the averaged embedding, our model can not only generate conformations close to the metastable conformations but also the pathway between them. We refer the above inference pipeline as “landscape explore” pipeline, whereas the standard pipeline uses the default Alphafold2 pipeline to extract embeddings. Fig 2 displays the schematic diagram of the techniques mentioned in this section.

#### 1. 3. MD simulations

The Amber22 program was used to perform molecular dynamics simulations in this study. High-resolution complex structures were obtained from PDB. For the simulation force field, CHARMM36 ^38^ was chosen, and the solvent water molecules were modeled using the TIP3 model. The nucleic acid molecules and proteins were initially positioned at the center of the water box in the simulation system. The solute molecules were kept at a distance of 20 Å from the surface of the water box. Additionally, 0.15 mM NaCl were added. The simulations were performed using periodic boundary conditions, with the temperature set to 300 K.

## Results

### 1. Protein hinge motion and ligand conformational selection

Protein structural changes can be groups into small shear motion and large hinge bending motion^39^. Shear motion refers to sliding movement of one structural region or domain on other parts of the tertiary structure. The F1-ATPase structures are good cases for the shear motion. Two inhibitors bind two shear motion conformers (TM-score 0.76), AlF3 in PDB 1E1R and BeF3 in 1W0J, respectively. In the current study, we generated conformers matches to both PDBs, with TM-scores of 0.92 (to 1e1r) and 0.93 (to 1W0J). Hinge motions involve relatively large position change of two domains caused by small local structural change of hinge linker, and a family of periplasmic sugar-binding proteins (pentose/hexose sugar receptors) involves the classic hinge motion from apo open state to ligand binding closed conformation^40^. We tested two of the periplasmic sugar-binding proteins D-allose binding protein and D-ribose binding protein. As can be seen in Figure 3, the Deepconformer generated conformations matching both and closed states with high accuracy.

**Fig 3.**
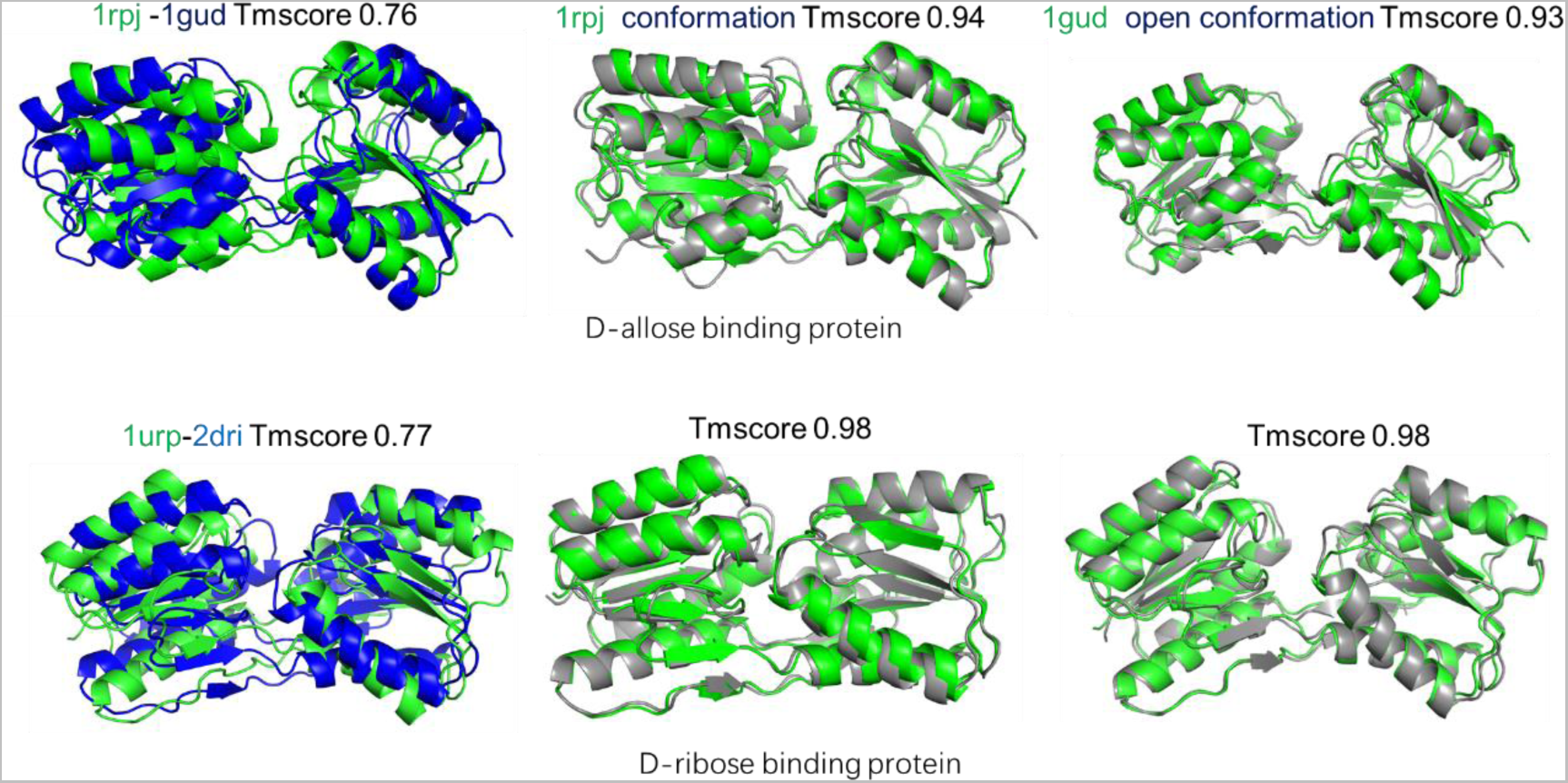
Generation of hinge motion conformers for D-allose binding protein (upper panel) and D-ribose binding protein (lower panel). Experimental structures were superimposed to show the structural difference between open and closed conformation (right). The superimpositions between the experimental structures (green) and their most similar (of the lowest RMSD) predicted structures (grey) were shown in middle (closed conformation) and right (open conformation).

Kinases often involve large conformation change during catalytic processes. Figure 4 compares two conformations of adenylate kinase, one with a transition state analog inhibitor AP5A (PDB: 1AKE), another is an apo state conformer. Two conformers differ by the large motion mixed with shear and hinge motions, with TM-score of 0.68 only. Again, Deepconformer recovered two conformations successfully, with TM-scores of 0.92 to closed 1AKE state (middle) and 0.94 to open state (4AKE, right).

**Fig 4.**
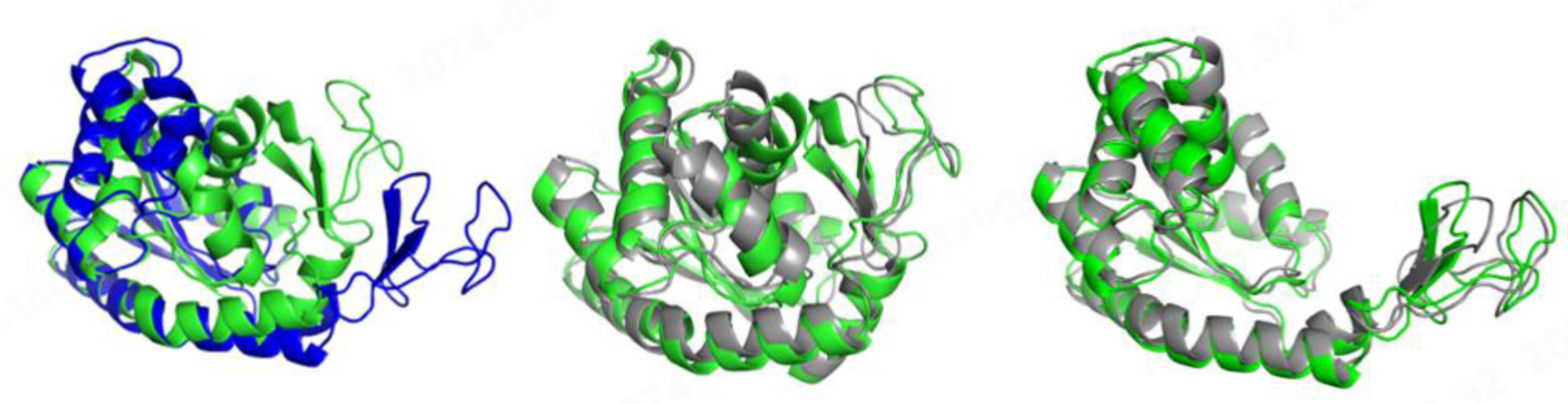
Catalytic conformation changes in Adenylate kinase

### 2. Fold Switch of KaiB protein

DeepConformer can generate distinct conformations of a natural protein and the corresponding conformations connecting their conformation transition pathway. KaiB is a circadian rhythm protein found in cyanobacteria and proteobacteria that adopts two conformations with distinct secondary structures (Figure 5): the “ground state” conformation which has a secondary structure of βαββααβ (PDB: 2QKE) and “fold-switch” (FS) conformation, which has a thioredoxin-like secondary structure (βαβαββα) (PDB: 5JYT). Standard AF2 pipeline predicts the FS state.

With our “landscape explore” pipeline, DeepConformer was able to generate conformation close to each native structure. We then locate intermediate pathway conformations by interpolating noise in the reverse process between the two conformations that are closest to two native structures. In Fig 6 we display two intermediate conformations together with experimental structure 2QKE and 5JYT.

**Fig 5.**
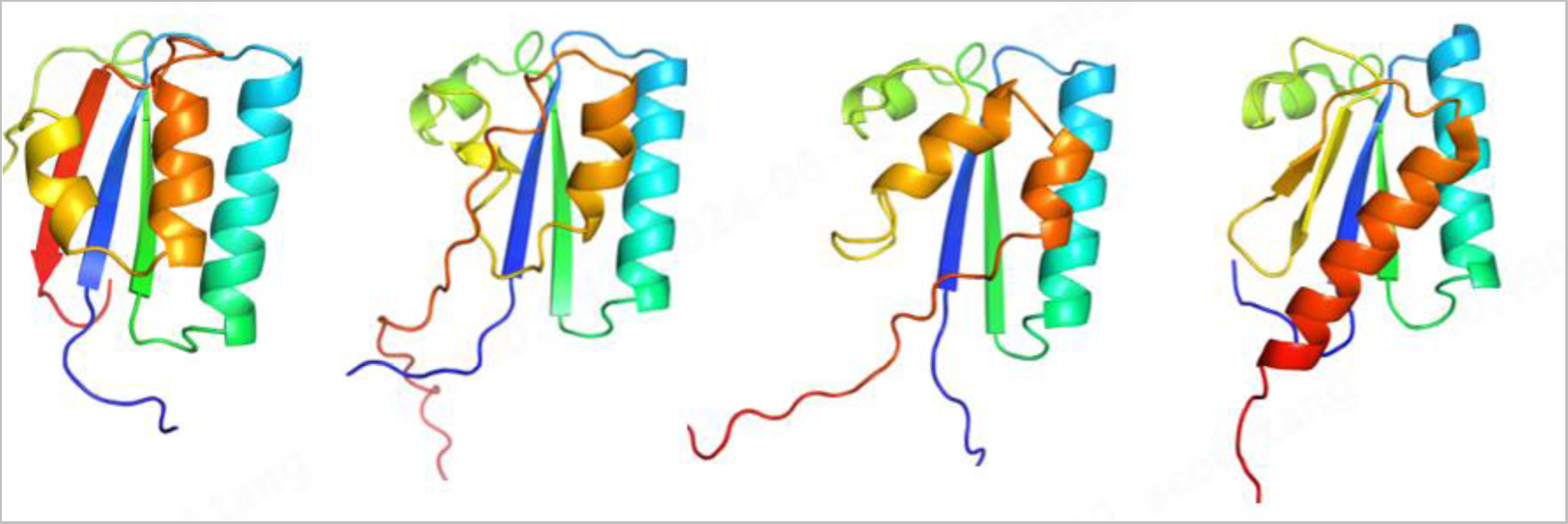
Fold switch of KaiB protein. From left to right are native structure 2QKE, intermediate structure A, intermediate structure B, native structure 5JYT. Rainbow color map is used to identify residue number. N-terminal corresponds to red color and C-terminal corresponds to blue color.

### 3. Dynamics features in DeepConformer generated protein conformations

Deepconformer does not use MD generated conformations for training. Here we examine Deepconformer generated protein conformations and compared with MD sampled conformations and dynamics. We first examined prediction results of our standard pipeline. We first used Adenosine A2A Receptor (PDB:3PWH) as an example protein. The DeepConformer-generated structures superimposed on experimental structure (PDB:3PWH) are displayed in Fig 5. The predicted conformations show good agreement with experimental structure. To verify whether DeepConformer-generated conformations capture dynamic property of target protein, we ran a 200 ns MD simulation for the same protein. We calculated the rooted-mean-square-fluctuation(rmsf) for both DeepConformer and MD sampled conformations (Figure 5). Though our training data does not include any MD simulation data, the rmsf of generated conformers show good agreement with MD results.

**Fig 5.**
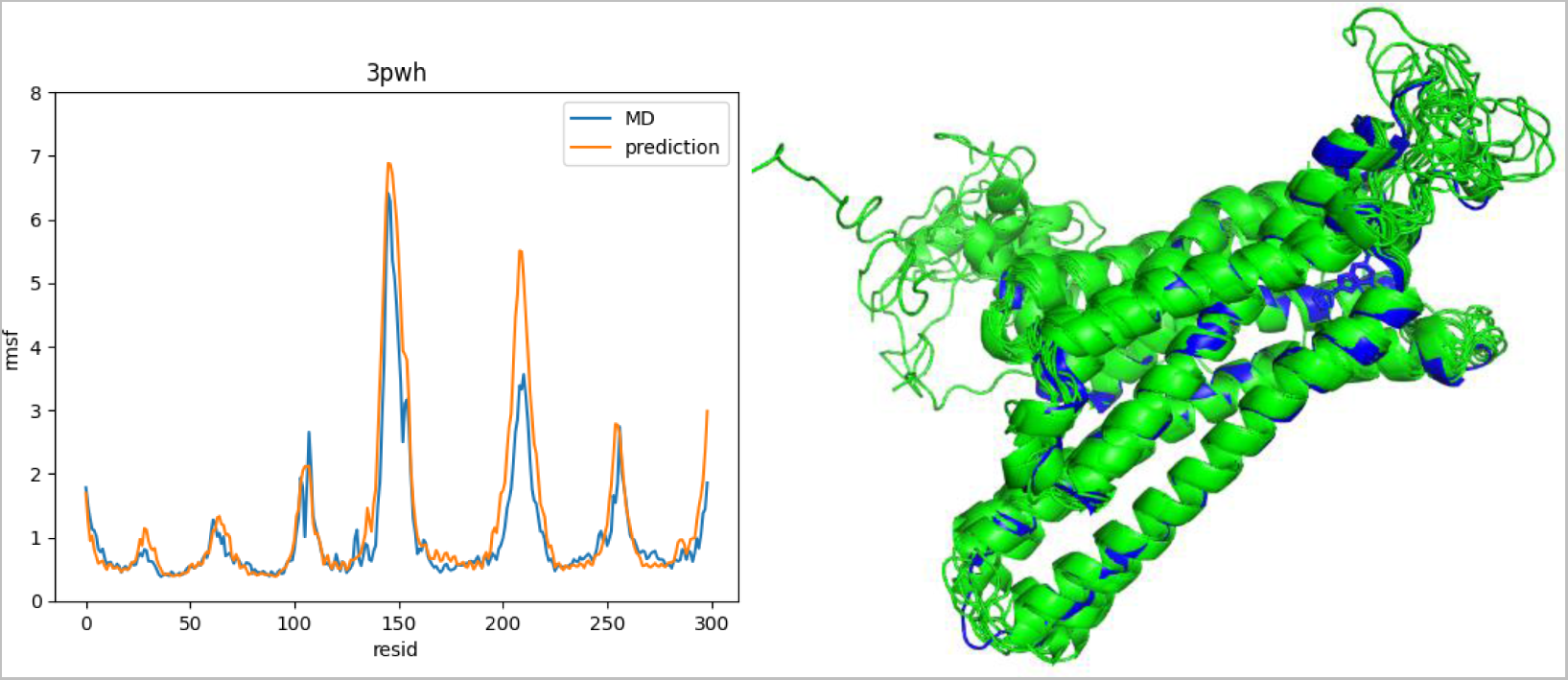
Left: RMSF of predicted conformations and that of MD conformations. Right: model generated conformations(green) superposed on PDB structure(blue) of Adenosine A2A Receptor.

We then used K-Ras protein (PDB 5kyk,5v71) as another example. Fig 6 displays the rmsf curve of DeepConformer-generated conformations and MD sampled conformations. Generally speaking, rmsf curve of model-generated conformations has good agreement with that of MD simulation. Though in the peak region (residue 25-40 and 60-75), the rmsf of model-generated conformations is lower than that of MD simulation. We further calculated the residue covariance matrix of model-generated and MD conformations and visualized in Figure 6. These two correlation matrixes show overall good agreement and the correlation coefficient of these two matrixes is 0.826. The agreements in both rmsf curve and covariance matrix with simulation data indicates that DeepConformer captured the underlying conformer dynamics.

Though conformations generated by DeepConformer with standard pipeline replicates small magnitude conformational flexibility well, it has difficulty to sample large magnitude conformation change. This is not surprising since our training data has very limited large magnitude conformational diversity and our model relies on embedding extracted from pretrained Alphafold2 model which may only contain information of one dominant conformation. To address this issue, we used the “landscape explore” pipeline introduced in previous section.

**Fig 6.**
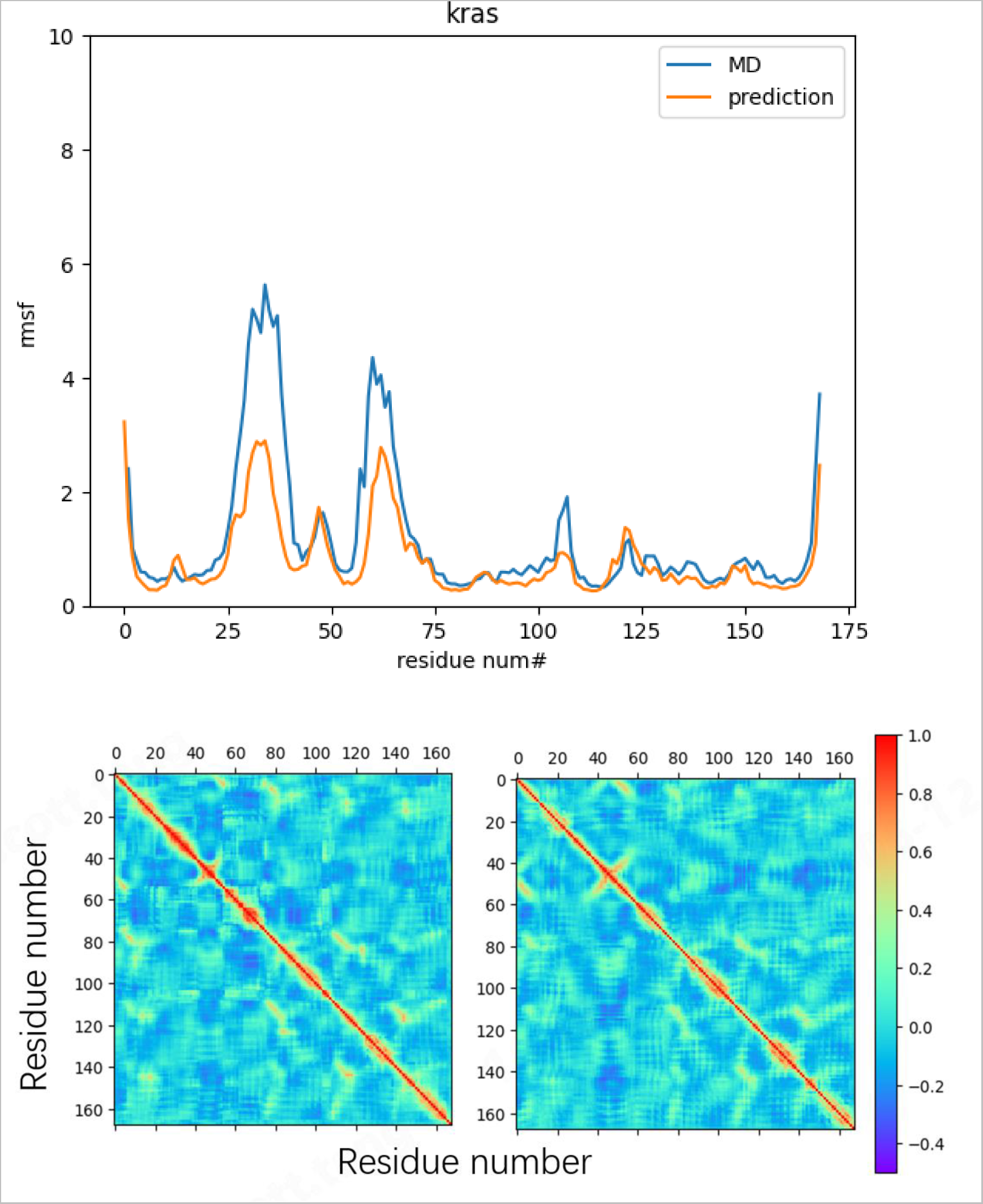
Up: RMSF of predicted conformations and that of MD conformations for Kras protein (pdb 5kyk). Down: Correlation matrix of MD (left) and prediction (right) conformations. The correlation coefficient of these two matrixes is 0.83.

The Kras protein has structural changes between different experimental structures. The most significant difference is the orientation of the helix and adjoining loop from residue 60 to 75. We visualized the DeepConformer-generated structures with both standard pipeline and the “landscape explore” pipeline on a two-dimensional plane (obtained with Multi-Dimensional Scaling or MDS^41^) together with the experimental structures (Fig 4). The conformations generated with standard pipeline are biased to PDB:4L9W and 4DSO while fail to cover the all other PDB structures. With the “landscape explore” pipeline the predicted conformation space is significantly enlarged and is able to cover all the PDB structures. In Fig 7 we also showed the 2 PDB structures and 2 generated structures, focus on the helix region from residue 65 to 75. DeepConformer with “landscape explore” pipeline was able to capture the major flexibility in this region. In table 1 we listed the smallest RMSD from each PDB structure to the predicted conformations as well as the largest RMSD from each PDB structure to the other PDB structures.

**Fig 7.**
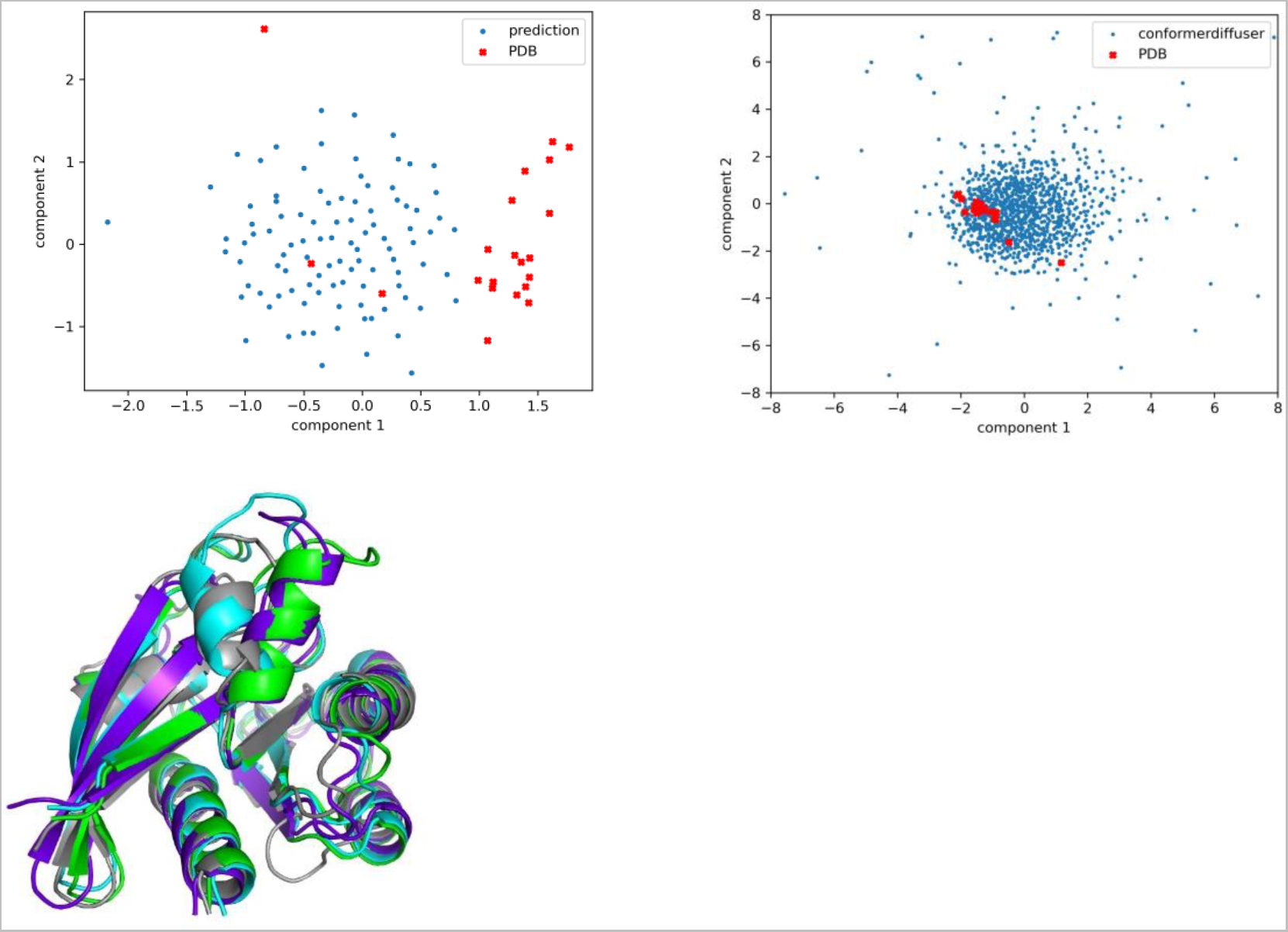
Up left: MDS plot of predicted conformations with standard pipeline and native structures from PDB. Up right: MDS plot of predicted conformations with standard pipeline and native structures from PDB. Down: blue and cyan denote two PDB structures of Kras protein. Grey and purple denote two generated conformations with lowest RMSD in the helix region from residue 65 to 75 from blue and cyan respectively.

**Table 1.** List of PDB structures covered in the Deepconformer generated conformations. Entries include the RMSDs of the experimental structures from the respectively most similar predicted structures (the smallest RMSDs) and from the most dissimilar (the largest RMSDs) PDB structures.

| PDB code | Min RMSD | Max RMSD |
| --- | --- | --- |
| 6h46 | 1.917 | 3.375 |
| 6p8w | 1.388 | 2.768 |
| 7c40 | 1.525 | 3.375 |
| 5xco | 1.454 | 3.093 |
| 5v9l | 1.479 | 3.214 |
| 7rt1 | 1.399 | 2.859 |
| 4l9w | 1.186 | 2.919 |
| 4lv6 | 1.322 | 2.708 |
| 8dni | 1.268 | 2.701 |
| 4q21 | 1.347 | 2.984 |
| 6n2k | 1.398 | 2.799 |
| 5v71 | 1.502 | 3.350 |
| 5e95 | 1.494 | 2.306 |
| 7u8h | 1.395 | 2.855 |
| 8afd | 1.427 | 2.840 |
| 5v9o | 1.489 | 2.909 |
| 6b0v | 1.470 | 2.862 |
| 7ewb | 1.398 | 2.863 |
| 5yxz | 1.430 | 3.052 |
| 6pgp | 1.358 | 2.780 |
| 4dso | 1.282 | 2.758 |

### 4. Comparison of MD simulations and Deepconformer generations of Domain motions of HIN-A and HIN-B in IFI16

IFI16 is an interferon-inducible DNA-sensor protein and plays important roles in antiviral response^42^ and aging^43^. IFI16 has two HIN domains, HIN-A from residues 199-389, and HIN-B from residues 520-705. Both HIN domains share very high structural similarity, each HIN domain has two subdomains which compact together and connected by a linker (Figure 8). The sequence identity between the HIN-A and HIN-B domains is around 40%, but their crystal structures (HIN-A: 2OQ0; HIN-B: 3B6Y) differ only by rmsd of 1.3 Å for all the main chain Cα atoms. The other HIN domains interact with p53 binding protein (p53bp) requires the conserved MFHATVAT motif (Figure 8) present in all 200 amino acid repeat regions of HIN-200 proteins^44^.

**Fig 8.**
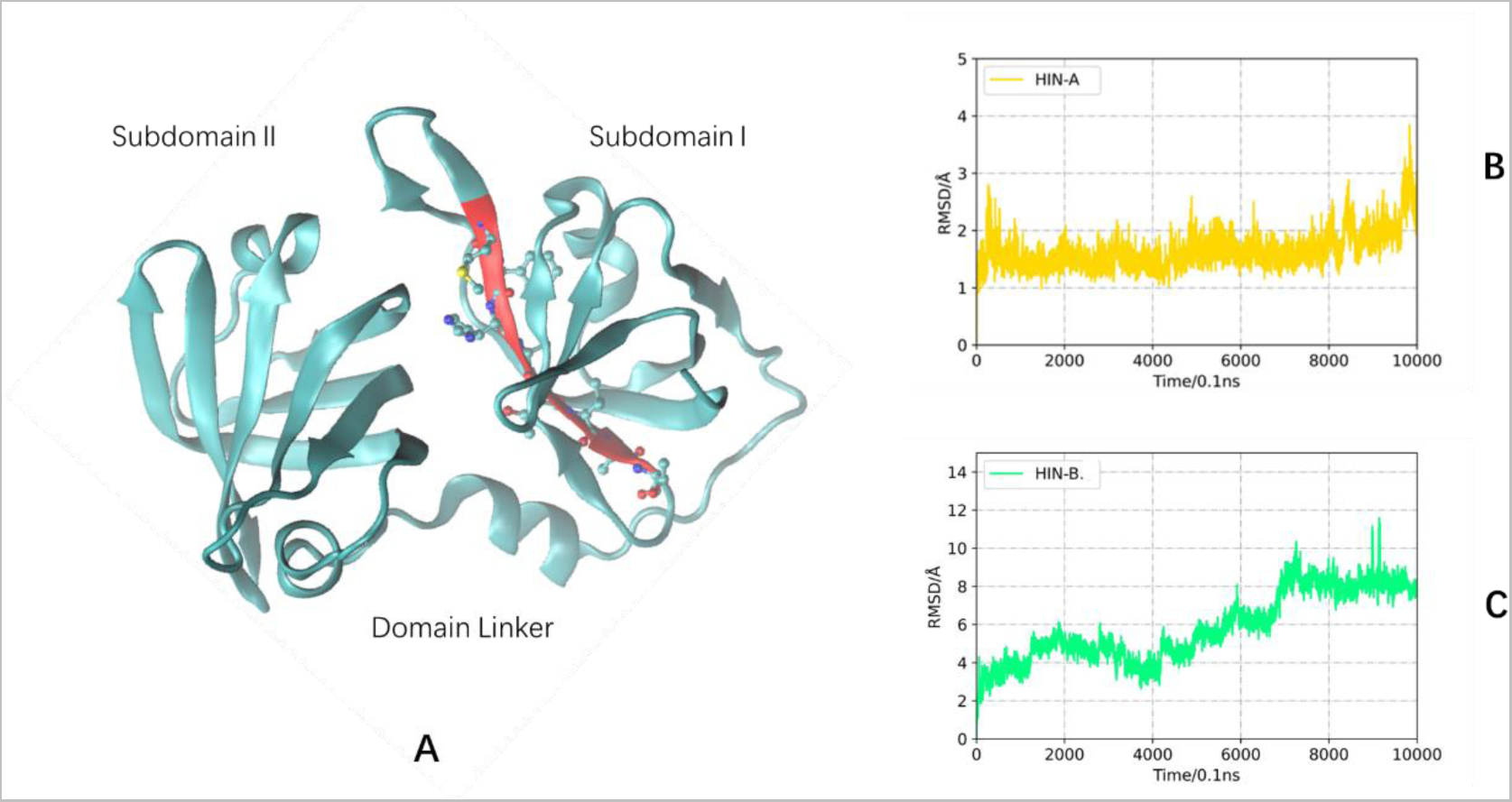
Domain structure and stability of HIN domains in IFI16. A: HIN domain has two subdomains which compact together and connected by a linker. Red ribbon is the MFHATVAT fragment which interacts with p53binding protein. B: RMSD trajectory indicates that HIN-A domain is stable during 1000 ns MD simulation. C: HIN-B domain shows flexibility during MD simulation.

The interactions of IFI16 and p53 not only affect p53 pathways, but also STING activation in response to DNA damage^45^ and other gene regulations^46^. Previous studies revealed that conserved MFHATVAT motif (Figure 8A) present in all HIN domains, but only HIN-A interact with p53^44^. Another study ^47^ also suggested that HIN-A domain binds to the basic C terminus of p53, whereas the HIN-B domain binds to the core DNA-binding region of p53. This study also suggested that p53 C-terminus binds to an overt acidic/hydrophobic patch observed on the surface of HIN-A (formed by residues Y218, T220, E222, Y267, E272, E381), but not HIN-B domain of IFI16. While the surface-patch binding site may explain differential selection of p53 C-terminus to HIN-A and HIN-B, the binding affinity of p53 C terminus binds IFI16 HIN-A domain was determined to be only Kd ∼20 μM.

The binding of MFHATVAT motif with p53bp require complete opening of subdomains of HIN-A and HIN-B, and p53 binds with different HIN domains in IFI16. These behaviors triggered us to study the domain motions of HIN-A and HIN-B using both MD simulations and Deepconformer conformational search. Interestingly, in the MD simulations, we found that the HIN-A is much stable than HIN-B. Two subdomains of HIN-A remain closely packed during 1000 ns simulation, indicated by steady state RMSD trajectory plot in Figure 8B. However, HIN-B already shows subdomain twisting and slight open, leading to a conformation trajectory with much larger RMSD (Figure 8C).

We used HIN-A to compare effectiveness of our dynamics enhancing techniques described in section 2.2. Fig 9 displays the predicted conformations of HINA(PDB 1oq0) with different models projected onto MDS plane. Model 1 is trained with PDB data without the embedding masking procedure, model 2 is trained with PDB expanded with swiss-model repository data without the embedding masking procedure, model 3 contains embedding masking procedure in training and inference process. Fig 9 indicates that both expand PDB data with swiss-model repository and randomly mask embedding effectively expand the distributional space of the predicted conformations. We then used Deepconformer generated 500 conformers for both HIN-A and HIN-B. Both HIN-A and HIN-B conformation ensemble contain structures with completely open subdomains (Figure 10A and 10B), indicating Deepconformer’s advantage over MD simulations to explore large conformation change. Figure 10C compares the Multidimensional scaling distributions of HIN-A conformers and HIN-B conformers. Near crystal structure region (red circle, distance < 5) HIN-A populated more conformers close to crystal structure than HIN-B, consistent with MD simulation results that HIN-A is more stable. However, the open conformations of HIN-A also has larger extent subdomain separation than HIN-B, as evident by large MDS distances for open conformers of HIN-A. Therefore, Deepconformer provided consistent conformation distributions with MD simulations in the local regions near crystal structures. Meantime, Deepconformer also generate large structural changes not readily accessible in MD simulations, and the large domain motions provided insights into biological function of IFI16.

**Fig 9.**
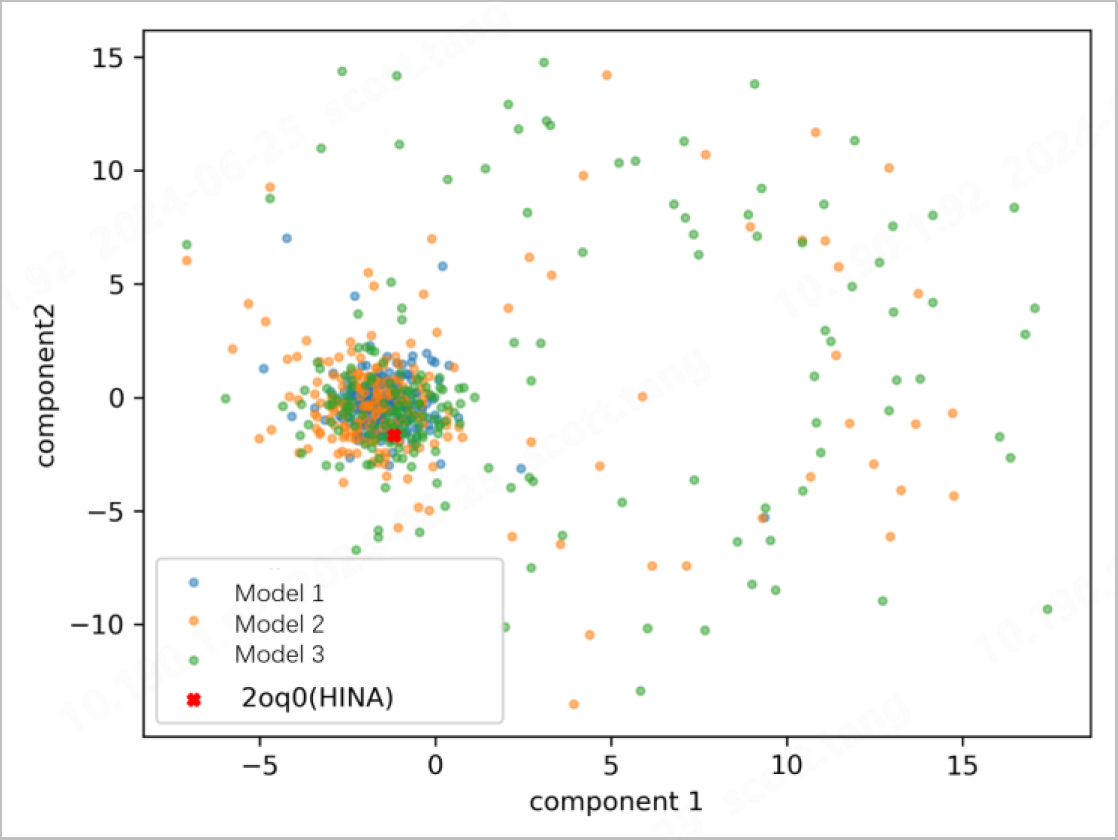
Predicted conformations of HINA with different models projected onto MDS plane.

**Fig 10.**
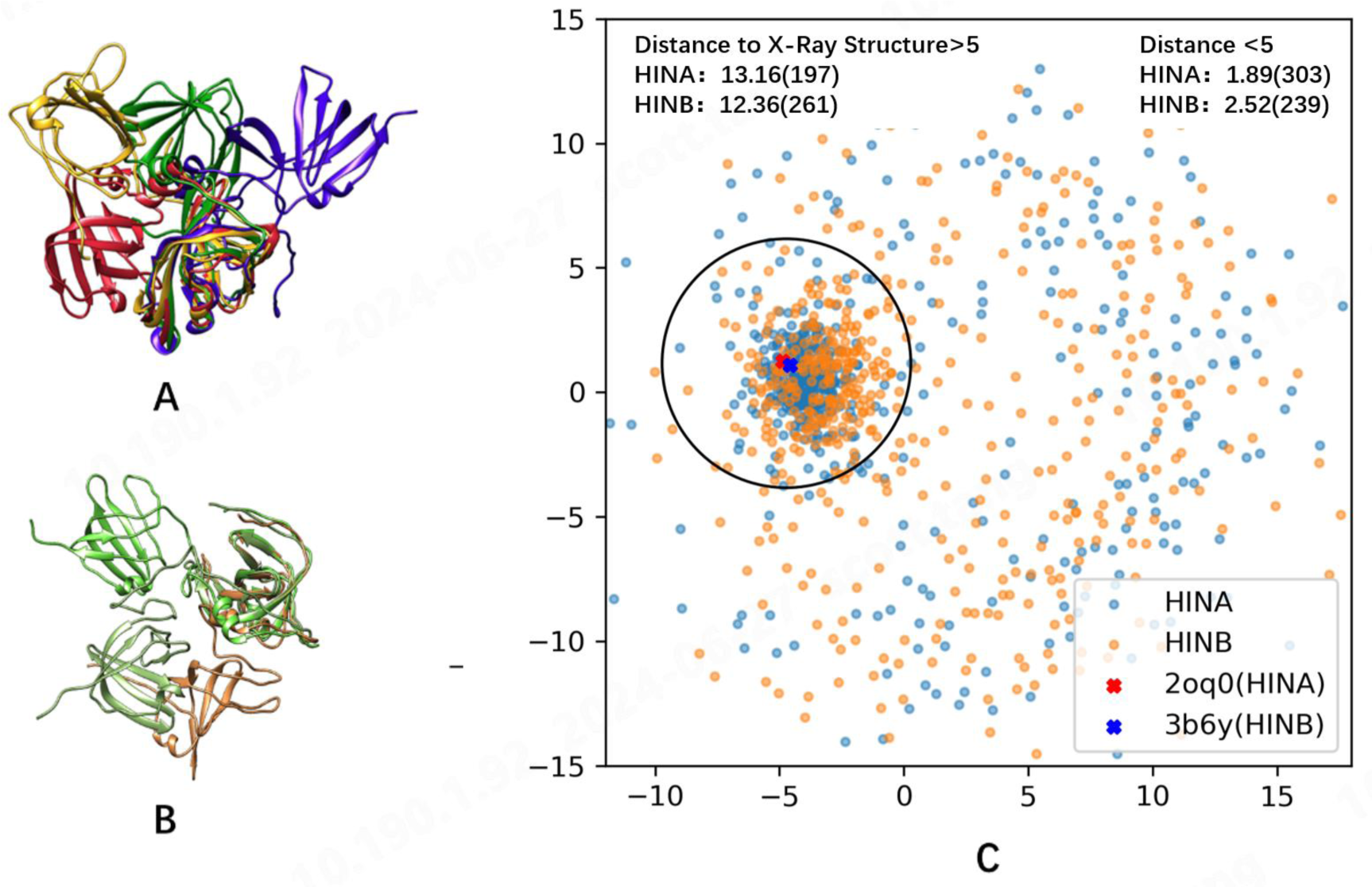
Conformational distributions of HIN-A and HIN-B in IFI16 generated by Deepconformer. A:Superimposing of open conformations of HIN-A on closed x-ray structure (red ribbon), B: Superimposing of three open conformations of HIN-B; C: Multidimensional scaling (MDS) plot of 100 conformers of HIN-A and HIN-B generated by Deepconformer.

## Discussion and Conclusion

Alphafold2 achieved breakthrough in protein structure prediction by unintentional embedding biophysical energy information from known protein structures, and the learned energy function has been used to rank the quality of candidate protein structures. In this work, we tested a hypothesis that protein energy landscape and conformational dynamics can be learned from experimental structures in PDB and coevolution data. We combined three approaches to allow deep learning techniques to extract hidden dynamics energy landscape information: expanded sequence-structure mapping, large scale 50% structure masking, and MSA clustering.

It has been proposed two decades ago that in solution proteins exist in a range of conformations, and biological protein interaction is better described as general conformational selection mechanism ^48^. Therefore, the exploration of conformational ensemble on protein energy landscape is more biologically relevant than identifying a signal structure^6,7^. Meantime, the rich structural variation and information for a given protein sequence and nearby mutants can be used to reliably extract protein conformational ensemble and dynamics features. Technically, we have shown that increasing structures associated with a given protein sequence combined with large scale amino acid masking greatly increased conformational diversity generated. Protein sequence evolution may correlate with structural dynamics^32^. Our clustering of MSA and subsequent average of the top ranked clusters correlated of protein conformational diversity and associated dynamics.

Overall, we have demonstrated the DeepConformer-generated structures has similar dynamic properties to that of MD simulation sampled structures and can cover distinct native structures of a single sequence. As DeepConformer achieved this without using protein-specific models or training with data like molecular dynamics trajectories, we expect it to be widely applicable for exploring the conformational dynamics of both natural and designed proteins to understand and optimize their function and regulation. One main limitation of this work is that we use Alphafold2 embeddings without finetuning. Information related with protein energy land scape from coevolution may be better retained in an end-to-end diffusion model. Another limitation is that physics regulations are not introduced in our model. Physics-based deep learning^49–51^ which can combine advantages of simulation and deep learning is a promising direction to solve protein energy landscape accurately and efficiently.

## Acknowledgments

B. Ma thanks support from Natural Science Foundation of China (Grant No. 32171246) and Shanghai municipal government science innovation grant 21JC1403700. This project is supported by Shanghai Digiwiser Biological, Inc.

